# Discovery of RNA polymerase α-subunit protein as a novel quorum sensing reprograming factor in *Pseudomonas aeruginosa*

**DOI:** 10.1101/2022.08.01.502315

**Authors:** Wenjie Cai, Huimin Liao, Mingqi Lu, Xiangting Zhou, Xiaoyan Cheng, Christian Staehelin, Weijun Dai

## Abstract

LasR is a master regulator of quorum-sensing (QS) in *Pseudomonas aeruginosa.* LasR-null mutants commonly appear in lung isolates from chronically infected cystic fibrosis (CF) patients. However, numerous such CF isolates have a QS-active phenotype, but factors underlying QS-reprogramming in LasR-null mutants remain largely unknown. Mutations in the transcriptional regulator gene *mexT* are well known to be responsible for QS-reprogramming in a laboratory LasR-null mutant strain, however, simultaneous occurrence of *lasR* and *mexT* mutations is rare in CF isolates. To identify QS-reprogramming determinants, we developed an experimental evolution approach, for which a QS-inactive LasR null mutant with an extra copy of *mexT* was engineered. In such a strain, spontaneous single *mexT* mutations are expected to have no or little phenotypic consequences. This novel method, named “targeted gene duplication followed by mutant screening” (TGD-MS), resulted in the identification of QS-active revertants with mutations in genes other than *mexT*. We characterized a QS-active revertant with a point mutation in *rpoA,* a gene encoding the α-subunit of RNA polymerase. QS activation in this mutant was found to be associated with the down-regulated expression of *mexEF-oprN* efflux pump genes. Our study therefore uncovers a new functional role for RpoA in regulating QS activity. Furthermore, our results suggest that a regulatory circuit controlling the expression of the *mexEF-oprN* operon is critical for QS-reprogramming. In conclusion, our study reports on the identification of non-MexT proteins associated with QS-reprogramming in a laboratory strain, shedding light on possible QS activation mechanisms in clinical *P. aeruginosa* isolates.

## Introduction

*Pseudomonas aeruginosa* is an opportunistic pathogen that causes severe acute and chronic human infections in cystic fibrosis (CF) patients with compromised immune system (1, 2). A cohort of *P. aeruginosa* virulence factors is under the control of the quorum-sensing (QS) system (3, 4). QS is a bacterial cell-cell communication system that regulates the expression levels of hundreds of genes in a cell density-dependent manner (4). *P. aeruginosa* has two acyl-homoserine lactone (AHL) QS systems, the Las and Rhl QS systems. In the Las QS system, LasI catalyzes the diffusible QS signal *N*-3-oxododecanoyl homoserine lactone (3OC12-HSL), while the transcriptional regulator LasR binds to the 3OC12-HSL forming a protein-signal complex. The active 3OC12-HSL-LasR complex further activates Las-regulon genes. Similarly, Rhl QS contains an RhlI-RhlR pair, with RhlI synthesizing the butyryl-homoserine lactone (C4-HSL) and the C4-HSL-bound RhlR activating expression of the Rhl-regulon (5). These two AHL QS systems further interact with a so-called *Pseudomonas* quinolone signal (PQS) system, which is responsible for generating 2-heptyl-3-hydroxy-4-quinolone (PQS) and its precursor 2-heptyl-4-quinolone (HHQ). The *pqs* genes (operon *pqsABCDE* and *pqsH*) code for HHQ and PQS synthesis (6, 7). In general, the Las QS is at the top of the QS hierarchy, and disruption of either LasI or LasR diminishes the expression of QS-activated genes (4).

QS-active *P. aeruginosa* show QS-regulated responses such as production of extracellular proteases, C4-HSL, hydrogen cyanide, rhamnolipids and pyocyanin (8–10). Secretion of QS-regulated proteases allow the bacteria to be visualized on agar plate containing skim milk (11, 12). Thus, spontaneous QS-active revertants derived from QS-inactive mutants can be selected on these agar plates. In such evolution experiments, mutants reprogramming their QS have been obtained from a QS-inactive *lasR* mutant of the laboratory strain PAO1 (13, 14). Isolates with mutated *lasR* gene have been identified in CF patients with chronic *P. aeruginosa* infection (8, 15, 16). However, although LasR is a master regulator of QS in *P. aeruginosa,* many LasR-null clinical isolates are QS-active (8–10). Hence, QS-reprogramming of LasR-null mutants also appears to occur under natural conditions.

The *P. aeruginosa* genome encodes several resistance-nodulation-division (RND) multidrug efflux pumps that transport small molecules into the extracellular environment. One of them, the MexEF-OprN efflux pump, is suggested to regulate the QS system by exporting the PQS precursor HHQ, which eventually leads to reduced intracellular PQS levels (17). Furthermore, MexEF-OprN can export antibiotics such as chloramphenicol, fluoroquinolones and trimethoprim, and thus contribute to antibiotic resistance (18). The *mexT* gene is localized upstream of the *mexEF-oprN* operon. The MexT protein, a LysR-type transcriptional regulator, positively regulates *mexEF-oprN* expression (19). In addition to the *mexEF-oprN* operon, MexT regulates expression of more than other 40 genes (20, 21). Expression of *mexT* is negatively regulated by its neighboring gene *mexS* (22, 23). Overexpression of the *mexEF-oprN* operon was found in various spontaneous *P. aeruginosa* mutants, and results in reduced QS-related responses, such as decreased synthesis of C4-HSL, hydrogen cyanide and pyocyanin (17, 24).

Mutations in the *mexT* gene were identified in clinical isolates from CF lungs and other patients with *P. aeruginosa* infections chronic CF lungs and other infections (15, 25) as well as in *P. aeruginosa* laboratory strains (26, 27). In a QS-inactive LasR-null mutant of the laboratory strain PAO1, spontaneous mutations in *mexT* were found to be sufficient for QS-reprogramming (13, 14), indicating that reduced expression of the *mexEF-oprN* genes is associated with a functional QS (20, 28). Likewise, disruption of the MexEF-OprN efflux pump in the LasR-null mutant background results in a QS-active phenotype with a functional Rhl QS as well as an active PQS system (13, 14). Deletion of genes of the PQS system (*pqsA* and/or *pqsE*) causes partially attenuated QS-dependent responses (13). Although mutations in *mexT* were frequently obtained in laboratory experiments, they are rare in clinical isolates (15). For example, a LasR-null isolate from a CF patient with chronic infection showed a QS-active phenotype while containing a functional MexT (9). Therefore, QS-reprogramming in such clinical isolates is expected to be controlled by factors other than MexT inactivation (non-*mexT* mutations).

In this study, we developed a novel experimental evolution approach to identify non-*mexT* determinants involved in QS-reprogramming. A QS-inactive *P. aeruginosa* LasR null mutant with an extra copy of *mexT* was generated and then used to screen for QS-active revertants. In this way, we identified a QS-active mutant with a spontaneous mutation in the transcriptional regulator gene *rpoA.* Further experiments showed that RpoA is involved in a regulatory circuit controlling the expression of the *mexEF-oprN* operon and thus critical for bacterial antibiotic resistance and QS-reprogramming. Hence, our developed mutant screening method not only resulted in the discovery of a novel function of the RNA polymerase α-subunit protein RpoA but also highlighted its association with the regulation of MexEF-OprN efflux pump activity.

## Results

### Simultaneous occurrence of *lasR* and *mexT* mutations is rare in *P. aeruginosa* CF isolates

LasR mutants are common in *P. aeruginosa* isolates from CF patients with chronic infection. These CF LasR-null isolates often contain an active Rhl QS (8, 10). In evolution experiments with QS-inactive LasR-null mutants of the laboratory strain PAO1, QS-active revertants were screened on skim milk agar plates and identified to have *mexT* mutations (13, 14). However, it is unclear whether such QS-reprogramming also applies to clinical isolates. To address this concern, we re-analyzed previously published whole genome sequencing (WGS) data of *P. aeruginosa* CF isolates with respect to *lasR* and *mexT* mutations (Smith et al., 2006). Among CF isolates from 29 patients examined, *lasR* mutations, including frameshift mutations, single nucleotide substitutions and insertions, were identified in isolates from 22 patients (75.9%). However, only isolates from two patients (9.1%) simultaneously had nonsynonymous mutations in *lasR* and *mexT* (Fisher’s exact test, *P* = 0.23, odds ratio = 0.27). Moreover, *mexT* mutations were additionally found in a number of isolates that contain an intact *lasR* (Table S1). Hence, *mexT* mutations are frequently not colocalized with *lasR* mutations in the examined WGS data. Therefore, inactivation of MexT might often not account for QS activity in clinical LasR-null isolates possessing a QS-active phenotype. In accordance with these findings, Cruz et al. (2020) reported on a QS-active clinical isolate which was identified as a LasR-null CF isolate with a functional *mexT* gene (9).

### Transposon mutagenesis results in a QS-active LasR null mutant with mutated *mexF* gene

As our analyzed WGS data and previous studies (9) both suggest non-MexT QS regulators are most likely responsible for QS-reprogramming in those LasR-null clinical isolates, we next sought out to identify such uncharacterized QS factors. We first performed a traditional transposon mutagenesis experiment using a *mariner*-based transposon (pBT20) (29) and a QS-inactive LasR null mutant of strain *P. aeruginosa* PAO1. The constructed insertion mutant library was spread and screened on skim milk agar. In this screening approach, QS-active revertant mutants were selected basing on the secretion of QS-controlled proteases. A total of 40, 000 mutant colonies, corresponding to a transposon density of about 156 bp per insertion in the PAO1 genome, were examined. Using this screening method, we obtained a single bacterial colony with a protease-positive phenotype. Sequencing analysis indicated that this mutant possesses a transposon insertion in the *mexF* gene, which codes for an essential component of the MexEF-OprN efflux pump. Given that QS-reprogramming in LasR null mutants mostly relies on mutations in *mexT* (13, 14), it was not surprising to see that a disruption of MexT-regulated MexF results in a QS-active phenotype. However, no other genes involved in QS-reprogramming were identified using this transposon mutagenesis method. Therefore, traditional transposon mutagenesis approach is not an efficient way to identify new QS-reprogramming factors in the LasR-null mutant.

### A novel experimental evolution method for identification of QS-reprogramming genes

Considering *P. aeruginosa* has an intriguing capacity for adapting to environments through frequent genome variations (30), we expected that hitherto unknown genetic elements involved in QS-reprogramming can be identified in evolution experiments. In a classic approach, a QS-inactive LasR-null mutant is evolved in casein medium and spontaneous mutants emerge. The QS-active phenotype of the protease-positive revertants can then be confirmed by measuring production of various QS-related metabolites such as pyocyanin. Using this approach, sequencing of QS-active mutants resulted only in mutations in the *mexT* gene (13, 14). We therefore developed a new approach in which the probability of obtaining *mexT* mutations was reduced by introducing an extra copy of *mexT* into a neutral site of the LasR mutant genome. We assumed that a deleterious mutation in one copy of the two *mexT* genes would still leave the other copy functional and therefore would have no or little phenotypic consequences in the evolution experiment. Using this strategy, mutants with single *mexT* mutations are expected to be largely filtered out, thereby facilitating the identification of non-*mexT* mutations. The method, termed “targeted gene duplication followed by mutant screening” (TGD-MS), is schematically illustrated in Fig. 1.

**Figure 1.**
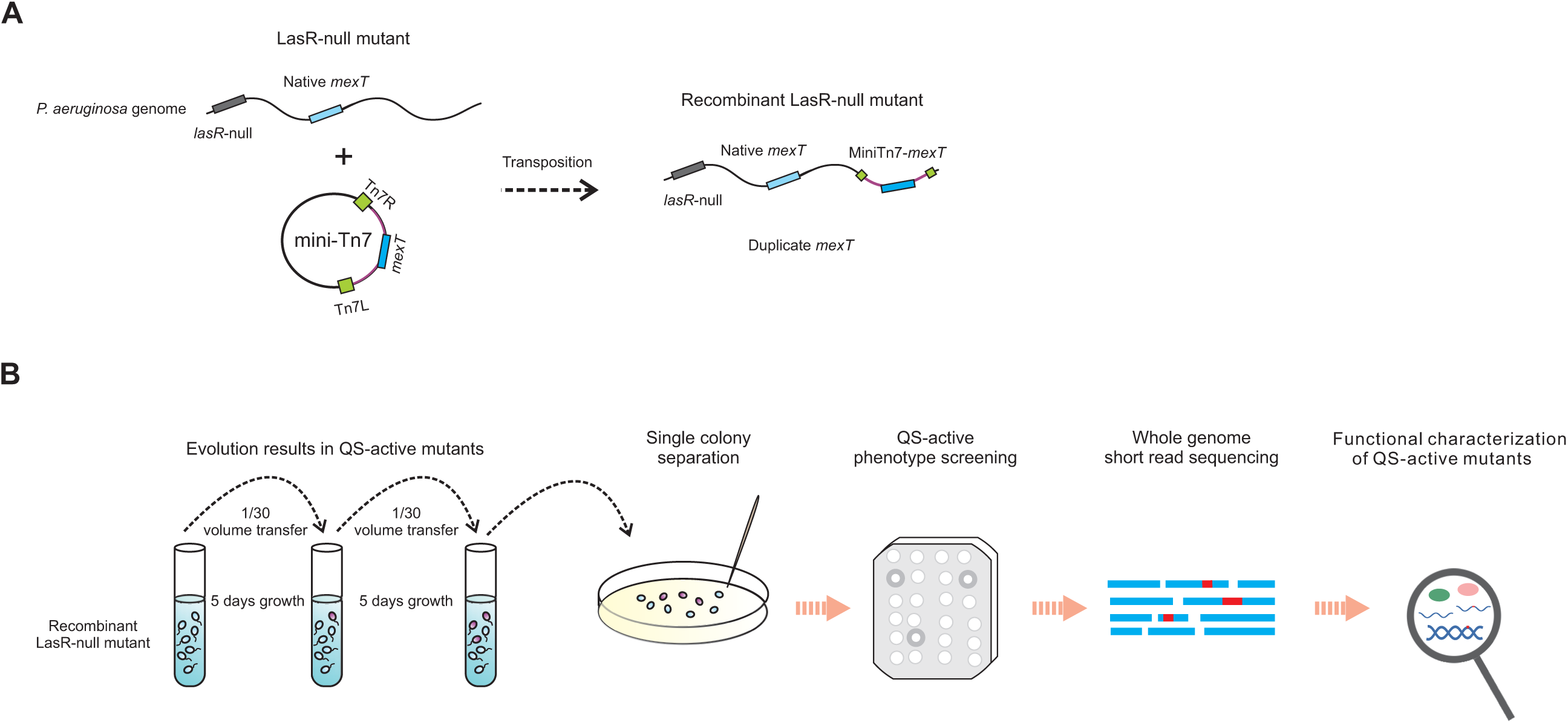
Scheme illustrating the novel TGD-MS strategy. (A) Illustration of the engineering of the recombinant LasR-null mutant. An extra copy of *mexT* was inserted into a neutral site of the LasR-null mutant genome, generating the recombinant LasR-null mutant carrying two copies of *mexT*. (B) The overall procedure of the TGD-MS strategy developed in our study aims to discover mutations in non-*mexT* genes. The evolution experiment is initiated with a QS-inactive recombinant LasR-null mutant with an extra copy of *mexT*. The bacteria are spread on skim milk agar plates to select QS-active revertants secreting QS-dependent proteases. This screening method results in the identification of mutations in genes other than *mexT*.

### The TGD-MS method results in the identification of a QS-active LasR null mutant with a mutation in *rpoA*

Using the established TGD-MS strategy, we screened for QS-active mutants of the engineered LasR-null mutant carrying two *mexT* copies. Mutants were further characterized with respect to the confirmation of QS-controlled extracellular protease activity using a skim milk agar assay and pyocyanin production. Four colonies were found to partially restore protease activity and show significantly increased pyocyanin production, providing clues that they contained a rewired active QS (Fig. 2). Three of these QS-active mutants were then subjected to whole-genome re-sequencing (WGS). The sequencing results revealed that these colonies possess a common single nucleotide mutation (nucleotide T4754641->C; amino acid residue T262->A) in *rpoA* (Table S2 and S3). The *rpoA* gene encodes the α-subunit of RNA polymerase (RNAP) in *P. aeruginosa*. Since *rpoA* is essential for cell viability in *P. aeruginosa* (31), a deletion mutant of *rpoA* has not been reported in the literature.

**Figure 2.**
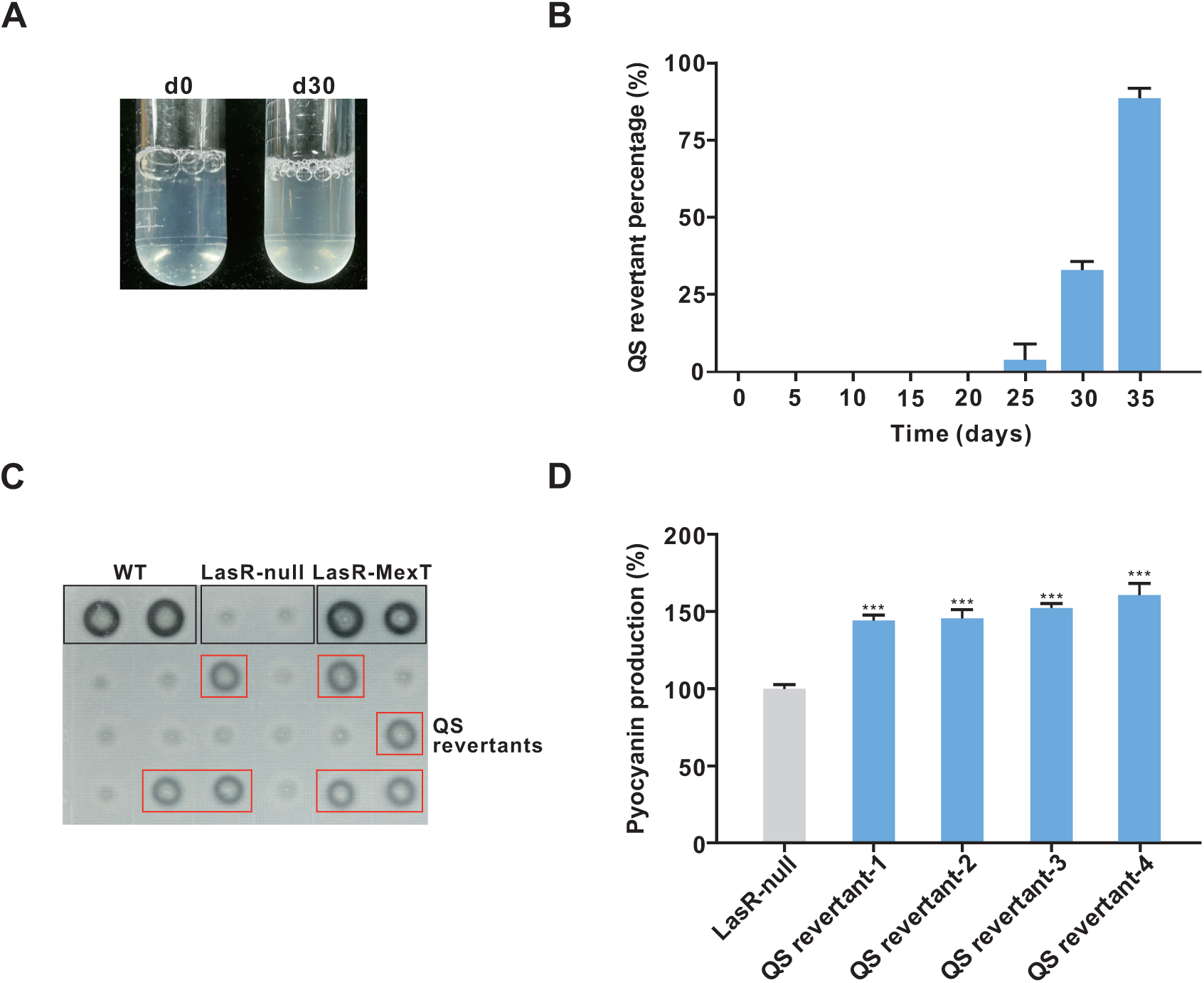
Identification of QS-active mutants obtained from a LasR mutant with an extra copy of *mexT*. (A) Photograph of culture tube of the LasR-null mutant carrying an extra copy of *mexT*, which was evolved at different time points in PM-casein medium. The culture of the engineered LasR-null mutant at d30 is slightly turbid as a result of QS revertants emergence. (B) Frequency of QS revertant colonies obtained from three TGD-MS experiments with the LasR mutant carrying an extra copy of *mexT*. QS revertant colonies were selected based on their protease-positive phenotypes on the skim milk agar plate. (C) Bacterial colonies were examined for extracellular protease activity at different time points using skim milk agar plates (control strains: WT, wild-type strain PAO1; LasR-null, LasR-null mutant used as a negative control; LasR-MexT, LasR-MexT employed as a positive control). Black rectangles indicate the control strains; Red rectangles show the QS revertants. (D) Pyocyanin production in identified protease-positive colonies. Pyocyanin production (OD_695_/OD_600_ values) was quantified for each strain (QS revertant-1/2/3/4, obtained mutants displaying protease-positive phenotype and significantly increased pyocyanin production). Data are presented as means ± SD (*n* = 3). \*\*\**P* < 0.001 (*t*-test).

To elucidate the role of *rpoA* in QS regulation, the same point mutation (T262->A) was introduced into the LasR-null mutant, generating the mutant LasR-RpoA-T262A (hereafter designated as LasR-RpoA* and the RpoA-T262A mutant as RpoA*). As expected, LasR-RpoA* displayed a typical QS-active phenotype with regard to extracellular protease activity as visualized on skim milk agar plates (Fig. S1). Furthermore, compared to the QS-inactive LasR-null mutant, LasR-RpoA* showed increased Rhl- and PQS-responsive activities as determined by reporter plasmids as well as elevated production of C4-HSL and PQS (Fig. 3). Overall, the QS-dependent activities and metabolites of the LasR-RpoA* mutant were higher than those of the LasR-null mutant, but lower compared to a constructed QS-active LasR-MexT double mutant (*mexT* deletion mutant of LasR-null). Complementation of the LasR-RpoA* mutant with an episomal copy of wild-type *rpoA* reduced the extracellular protease activity and pyocyanin production to levels determined for the LasR-null mutant carrying the empty vector (Fig. S2). In conclusion, application of the TGD-MS method resulted in the identification of RpoA as a novel regulator of QS-controlled processes, capable of rewiring QS activity in LasR-null.

**Figure 3.**
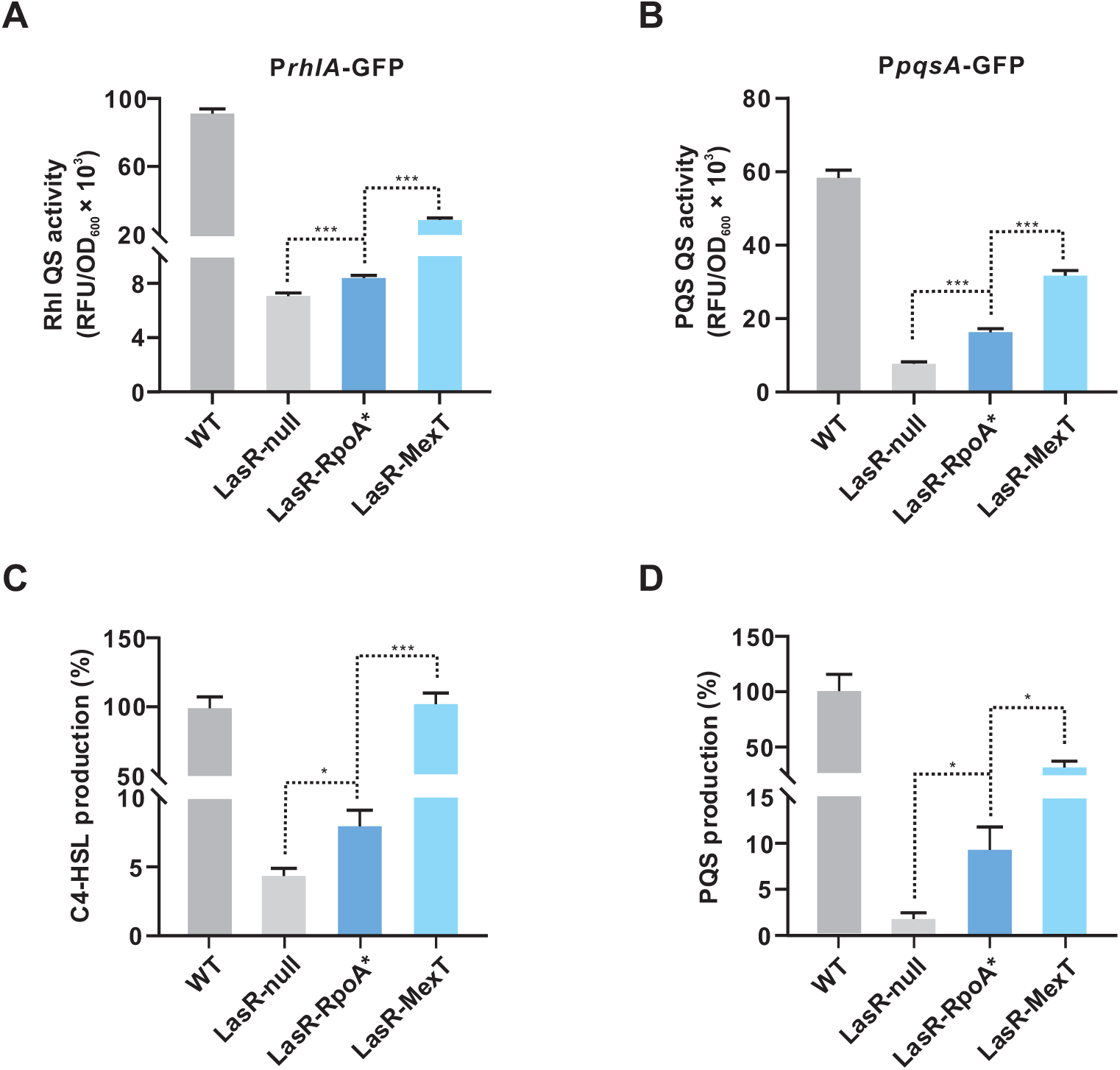
The LasR-RpoA* mutant shows increased QS-related activities. (A-B) Rhl- (A) and PQS-responsive (B) QS activities of the indicated strains. Rhl- and PQS-responsive QS activities are reflected by the fluorescence levels of the expressed reporters P*rhl*Α*-*GFP and P*pqsA*-GFP, respectively. Fluorescence values, expressed as relative fluorescence units (RFU), were obtained from bacteria cultured for 18 h. (C-D) the relative concentrations of C4-HSL(C) and PQS (D) in the shown strains. Production of AHLs in wild-type (WT) PAO1 was set to 100%. Data are presented as means ± SD (*n* = 3, *t*-test). \**P* < 0.05, \*\**P* < 0.01, \*\*\**P* < 0.001 (*t*-test).

### The LasR-RpoA* mutant exhibits a QS-active transcriptome profile

To investigate genome-wide gene expression changes in the LasR-RpoA* mutant, we performed an RNA-sequencing (RNA-seq) analysis and compared the transcriptome profile of LasR-RpoA* with that of the parent strain LasR-null. We found that a global gene expression was induced in the LasR-RpoA* mutant, with a total of 182 genes differentially expressed (at least a 2-fold change, *P* < 0.05) (Fig. 4A and Table S4). These differentially expressed genes were enriched in five pathways as presented by KEGG mapping (Fig. 4B). The QS circuit was among those enriched pathways, particularly with an upregulated *rhlI* gene and a strong induced *pqs* operon genes required for PQS production (Fig. 4C and Table S4). This transcriptome analysis confirmed that LasR-RpoA* possesses an active Rhl and a PQS QS systems. Because reduced expression of *mexEF-oprN* operon genes caused by *mexT* mutations resulted in QS-reprogramming in the LasR-null background (13, 14), we examined expression changes of *mexEF-oprN* operon genes in the LasR-RpoA* mutant. Likewise, LasR-RpoA* showed substantially decreased expression of genes encoding the MexEF-OprN efflux pump (Fig. 4D and Table S4). Of particular concern, no significantly expressed genes were observed in other 11 efflux pumps encoded in *P. aeruginosa* genome, indicating a specific downregulation of MexEF-OprN efflux pump in the LasR-RpoA* mutant. Finally, quantitative real-time PCR (qRT-PCR) analysis confirmed the data obtained by RNA-seq with selected QS-related genes and *mexEF-oprN* operon genes (Fig. S3).

**Figure 4.**
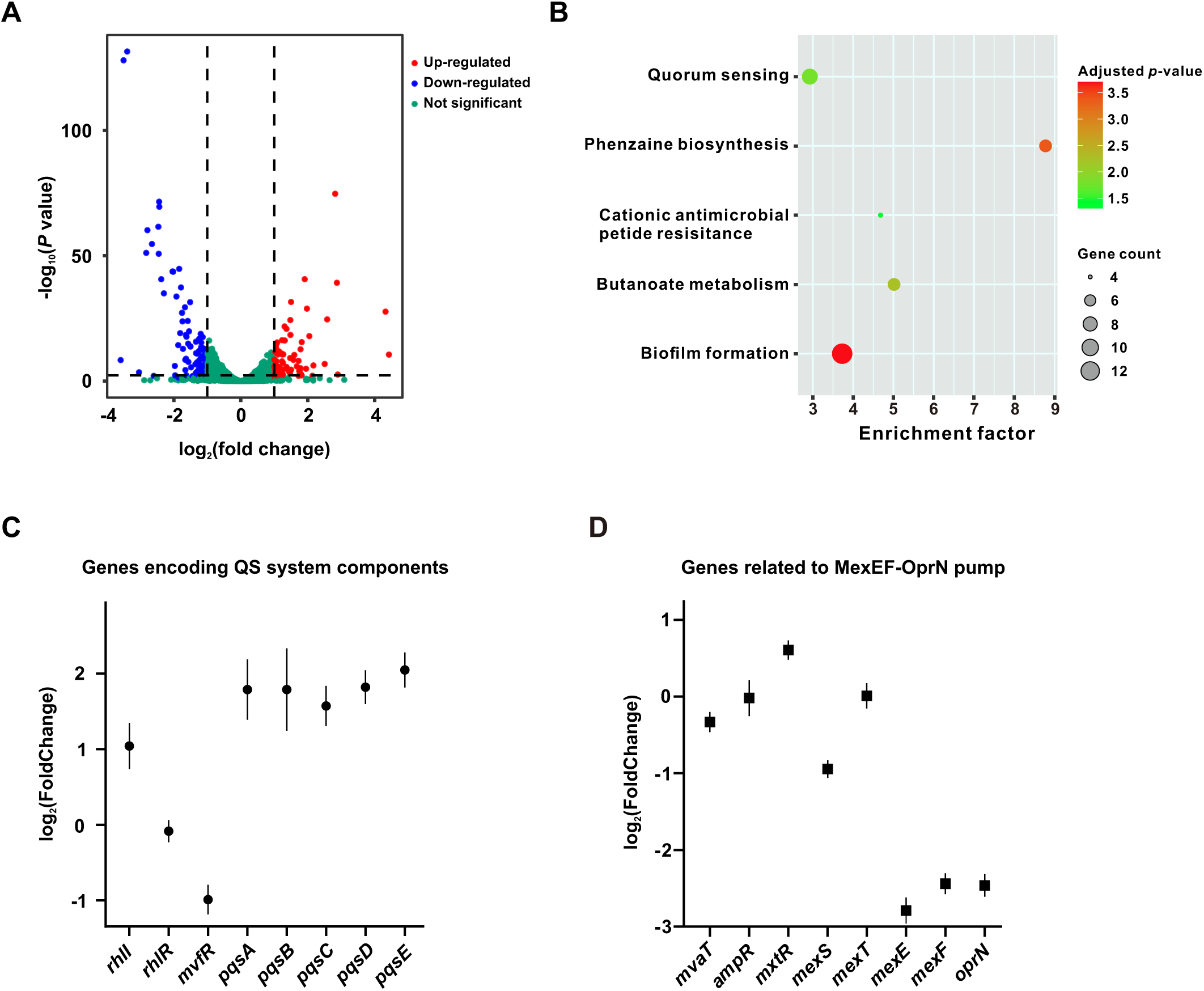
Transcriptome analysis of the LasR-RpoA* mutant. (A) Volcano plots showing the magnitude of differential gene expression induced by the identified T262->A mutation in *rpoA* (transcriptome comparison of LasR-RpoA* compared to the parent strain LasR-null). Each dot represents one annotated sequence with detectable expression (red dots, up-regulated genes; blue dots, down-regulated genes; green dots, genes that are not differentially expressed). Thresholds for defining a gene significantly differentially expressed (log_2_ (fold change) ≧ |1.0|, *P* value ≦0.05) are shown as dashed lines. (B) KEGG pathway analysis of differentially expressed genes. Fold enrichment shows the enrichment of differentially expressed genes in the corresponding pathway (differentially expressed gene number to the total gene number in a certain pathway). The size and color of the bubble represent the amount of differentially expressed genes enriched in the pathway and enrichment significance, respectively. (C-D) Expression levels of genes encoding QS system components (C) and MexEF-OprN efflux pump and its regulators (D). A positive log_2_ (fold change) value indicates up-regulation and a negative value indicates down-regulation of a given gene in the LasR-RpoA* mutant compared to LasR-null. Data are indicated by means ± SD.

Based on the transcriptome profile obtained in this study, we hypothesized that the attenuated MexEF-OprN efflux pump might account for the Qs-reprogramming observed in the LasR-RpoA* mutant. To investigate this speculation, we conducted a comparative transcriptome analysis between the LasR-RpoA* mutant and the LasR-null mutant. Our transcriptome analysis revealed that a total of 297 differentially expressed genes were induced in the LasR-MexT mutant (Table S5). Consistent with previous findings (21), many of these induced genes were associated with the MexEF-OprN efflux pump in the LasR-MexT mutant. Since the genes substantially upregulated by MexT inactivation are known to be MexEF-OprN-dependent (21), we selected genes with at least a 4-fold change in expression in the LasR-Mext mutant for further analysis of expression shifts in the LasR-RpoA* mutant. The comparative transcriptome analysis demonstrated that genes significantly upregulated in the LasR-MexT mutant were also activated in the LasR-RpoA* mutant (*P* = 0.01, one-sided Kolmogorov-Smirnov (KS) test). However, we did not observe a similar transcription pattern for down-regulated gene sets between the LasR-MexT mutant and the LasR-RpoA* mutant (*P* = 0.07, KS test). Given that disruption of the MexEF-OprN efflux pump is known to be responsible for QS-reprogramming in the LasR-MexT mutant (13, 14), we reasoned that similar to LasR-MexT, the reduced expression of *mexEF-oprN* operon genes might also be associated with QS-reprogramming in LasR-RpoA*.

### The RpoA* protein modulates the transcription of *mexEF-oprN* operon genes

We used a green fluorescent protein (GFP) based transcriptional fusion system (P*mexE*-GFP) to estimate the promoter activity of *mexEF* in PAO1 derivatives (core promoter of the *mexEF-oprN* operon). In line with the results obtained from RNA-seq and qRT-PCR analysis, the LasR-RpoA* mutant carrying the reporter construct caused a sharp reduction of fluorescent signals when compared to the LasR-null mutant, indicating the reduced expression of *mexEF-oprN* operon genes (Fig. 5A). An even stronger reduction of fluorescent signals was observed for the LasR-MexT mutant carrying the reporter construct (Fig. 5A). These findings are in agreement with the observation that QS-dependent activities are weaker in LasR-RpoA* than in LasR-MexT (Fig. 3).

**Figure 5.**
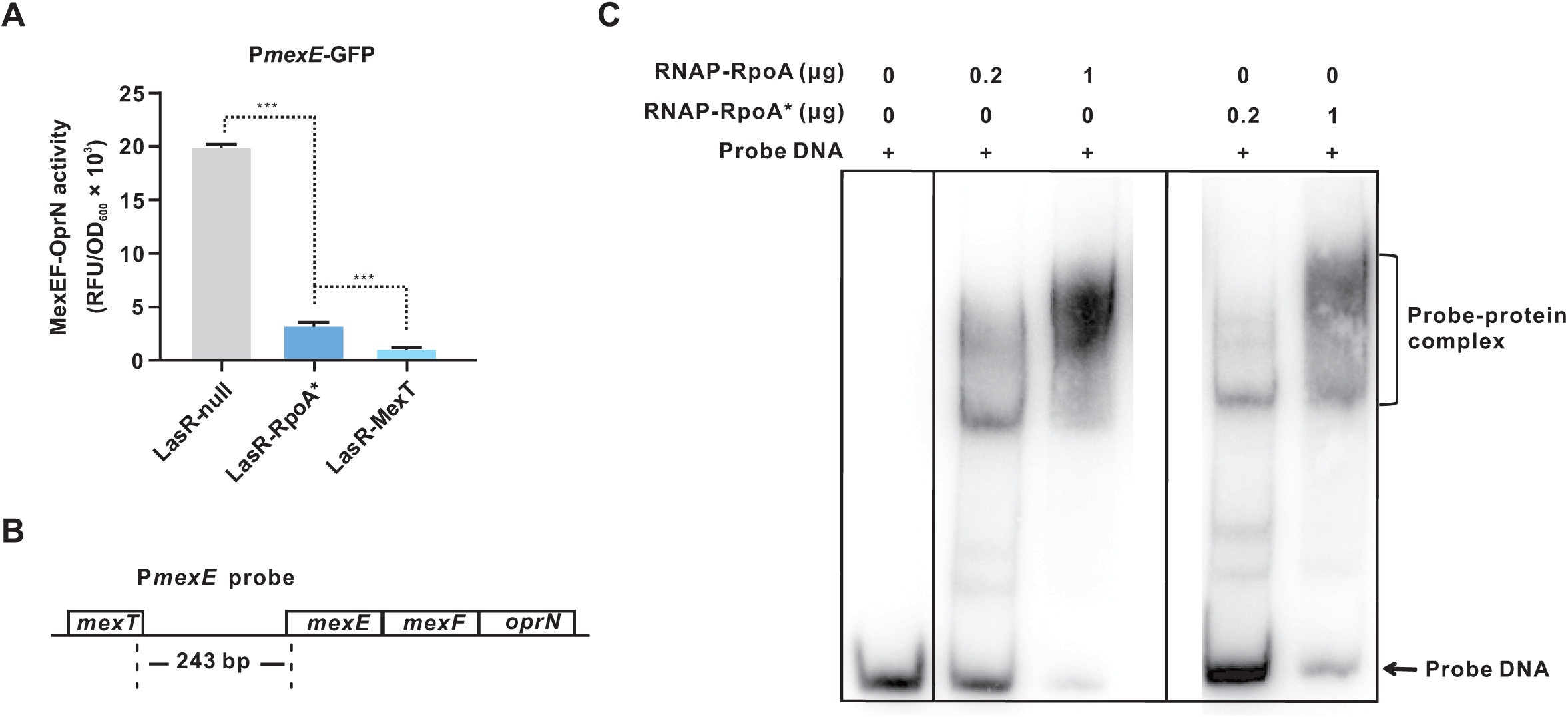
RpoA* modulates expression of *mexEF-oprN* operon genes. (A) The LasR-RpoA* mutant shows reduced transcription of *mexEF-oprN* operon genes as estimated with the help of P*mexE*-GFP, a *mexEF* promoter-GFP fusion construct. Data of indicated strains are presented as means ± SD. \**P* < 0.05, \*\**P* < 0.01, \*\*\**P* < 0.001 (*t*-test). (B) Illustration of the probe DNA used for the performed EMSA with recombinant RpoA of wild-type (WT) bacteria or RpoA*. (C) EMSA results showing that RNAP containing RpoA forms a DNΑ-protein complex. A weaker signal is seen for the sample with recombinant RpoA*. The unbound probe DNA is seen at the bottom of the blot.

We next examined how RpoA* attenuates the expression of *mexEF-oprN* operon genes. RpoA protein comprises α-NTD and α-CTD two domains, forming a core unit of the RNAP (32). Given the RpoA* mutation localizes in the α-CTD, a domain responsible for the contact of RNAP to promoter DNA, we hypothesized that the RpoA* protein may directly interfere with the binding of RNAP to the *mexEF* promoter. To test this hypothesis, we performed an electrophoretic mobility shift assay (EMSA). His-tagged RpoA or RpoA* was expressed in the LasR-RpoA* mutant. The recombinant RNAP proteins containing the His-tagged α-subunit were then extracted and purified by affinity chromatography. The two different types of RNAP were then assayed for their ability to bind to a 243-bp *mexEF* promoter DNA probe (Fig. 5B). Both RNAP proteins bound to the test probe in the EMSA (Fig. 5C). Compared to RNAP containing the wild-type RpoA, however, a weaker signal was observed for RNAP containing the RpoA* variant, which indicates a reduced RNAP-probe interaction caused by RpoA*. We therefore concluded that RNAP containing RpoA* down-regulates the expression of *mexEF-oprN* operon genes by reducing the direct contact of RNAP to the *mexEF* promoter DNA.

Previous findings showed that inactivation of the MexEF-OprN efflux pump up-regulates QS-dependent activities partially by reducing the export of the PQS precursor HHQ (17, 33), which may contribute to increased PQS production in LasR-RpoA* and LasR-MexT mutants (Fig. 3). As *pqsA* codes for a coenzyme ligase essential for PQS biosynthesis (34), we deleted the *pqsA* gene in the LasR-RpoA* mutant (generating LasR-RpoA*-PqsA) and evaluated the mutant’s ability to secrete proteases. The *pqsA* deletion caused a strong but not complete reduction of the QS-controlled extracellular protease activity (Fig. S4). Consistent with the observation of *pqsA* deletion in LasR-MexT (13), our obtained results suggest that QS-reprogramming in the LasR-RpoA* mutant also partially depends on increased PQS production.

Furthermore, we complemented the LasR-RpoA* mutant by transferring an episomal copy of the *mexEF-oprN* operon fragment driven by an *rrnB* promoter. Expression of *mexEF-oprN* operon genes restored QS-controlled pyocyanin production (Fig. S5), confirming the reduction of MexEF-OprN activity responsible for QS-reprogramming in LasR-RpoA*. In addition, we analyzed the LasR-RpoA* mutant with respect to swarming motility, as inactivation of the MexEF-OprN efflux pump was reported to stimulate swarming motility (33). As expected, the LasR-RpoA* was found to exhibit considerably increased swarming motility when compared to the LasR-null parent strain (Fig. S6).

### The LasR-RpoA* mutant is fitter than the LasR-MexT mutant in a casein medium supplemented with chloramphenicol

Given that a functional MexEF-OprN efflux pump promotes antibiotic resistance (24, 35), LasR-RpoA* thereby may reduce antibiotic resistance by attenuating the activity of the MexEF-OprN efflux pump. Due to the intermediate expression level of *mexEF-oprN* operon genes in LasR-RpoA* when compared with LasR-null and LasR-MexT, we assumed that LasR-RpoA* may also contain moderate antibiotic susceptibility. As a test of our assumption, we sought to investigate the resistance of LasR-RpoA* to chloramphenicol. Chloramphenicol is a known antibiotic substrate of the MexEF-OprN efflux pump (36). In comparison to LasR-null, LasR-RpoA* expectedly decreased resistance to chloramphenicol, while LasR-MexT showed higher susceptibility (Fig. 6A).

**Figure 6.**
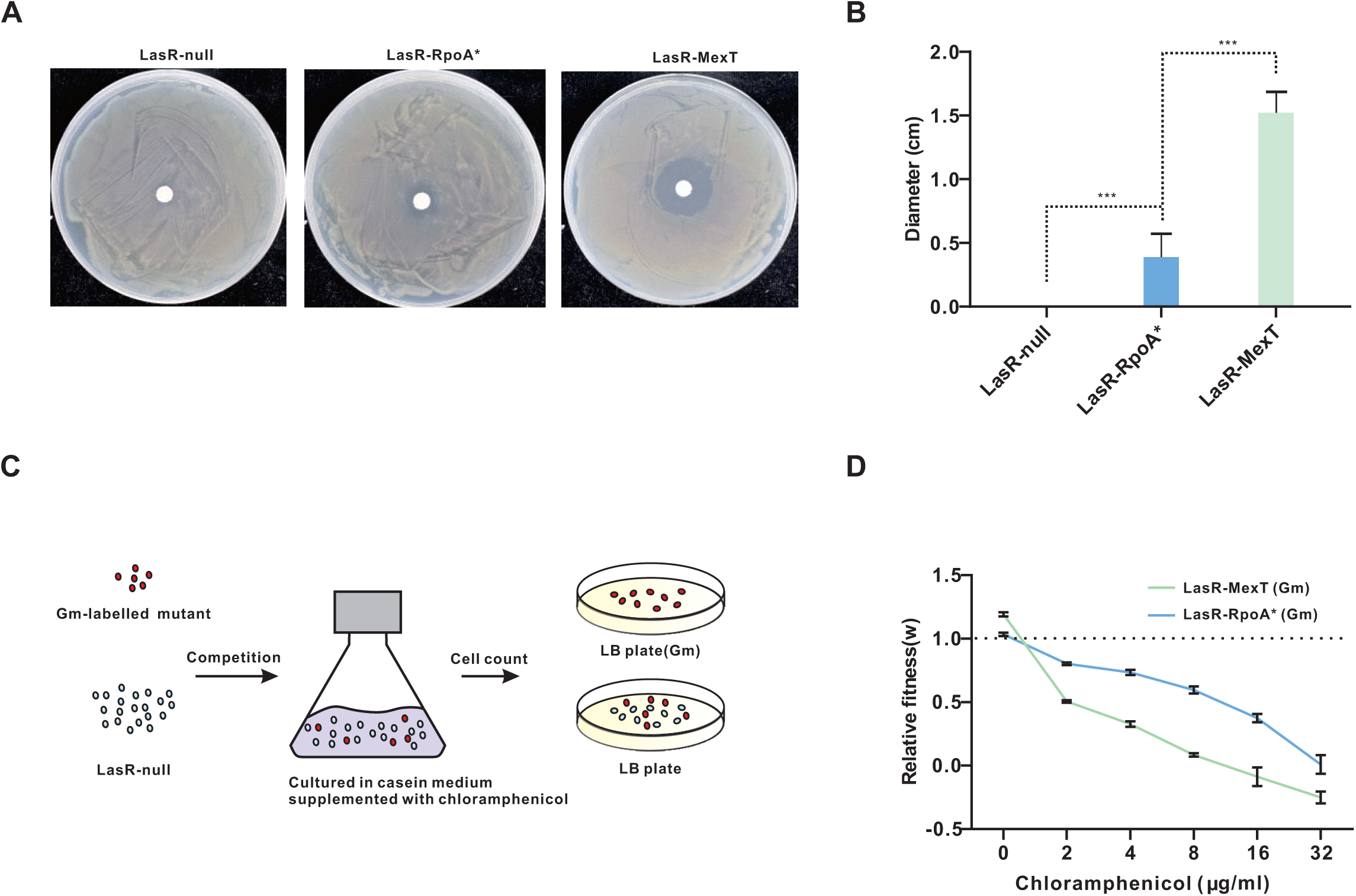
Effects of chloramphenicol on the growth fitness of the LasR-RpoA* mutant. (A)Sensitivity of the indicated mutants to chloramphenicol. The estimated 10^5^-10^6^ CFU of each tested strain was spread onto LB plates containing paper disks loaded with 40 μg/ml chloramphenicol. The photograph was taken after 24 h of incubation at 37°C. (B) Quantification of diffusion diameters formed around the paper disk in (A). (C) Experimental set-up of the competition assay. The LasR-RopA* and LasR-MexT mutants were labeled with a Gm resistance gene (pUC18T-mini-Tn7T-Gm). Each mutant was then co-cultured with the LasR-null mutant at a start ratio of 5:95 in PM-casein medium containing the indicated concentrations of chloramphenicol. After incubation for 48 h, colonies were counted and relative fitness (*w*) was estimated by determining the ratio of Malthusian growth parameters. (D) Determination of the relative fitness of LasR-RpoA* or LasR-MexT against LasR-null using the competition assay.

Through the regulation of the MexEF-OprN efflux pump, LasR-RpoA* thus gained a partially restored QS activity but a slightly reduced antibiotic resistance. We hypothesized that the arising LasR-RpoA* mutant from the neighboring LasR-null mutant would be fitter than the LasR-MexT mutant when the bacteria are exposed to an environment required both QS activation and antibiotic resistance. To test this hypothesis, we examined the ability of LasR-RpoA* or LasR-MexT to compete with LasR-null using the casein medium broth containing different concentrations of chloramphenicol. As expected, in the absence of chloramphenicol, LasR-MexT or LasR-RpoA* outcompeted LasR-null in the casein medium (Fig. 6B), consistent with the observation that QS-active LasR-null mutants are advantageous in the casein environment as previously reported (13, 14). In the presence of chloramphenicol, growth of LasR-RpoA* relative to LasR-null was reduced with increasing concentrations of chloramphenicol, indicating that chloramphenicol represses the growth advantage of LasR-RpoA*. Similarly, increasing chloramphenicol concentrations caused strongly reduced growth fitness of LasR-MexT relative to LasR-null. Notably, growth fitness reduction of LasR-MexT was even more pronounced than that of LasR-RpoA* (Fig. 6D). In conclusion, our findings indicate that the LasR-RpoA* is relatively fitter than the LasR-MexT mutant once it emerges from the LasR-null parent strain in the presence of chloramphenicol.

### Trade-off between QS-reprogramming and bacterial pathogenesis in LasR-RpoA*

To investigate the impacts of LasR-RpoA* on bacterial pathogenesis, we first evaluated the QS-controlled virulence products in the LasR-RpoA* mutant. As shown in Fig. S7, the LasR-RpoA* mutant exhibited higher levels of pyocyanin and hydrogen cyanide production compared to the LasR-null mutant, albeit lower than the LasR-MexT mutant. This intermediate level of QS-controlled virulence factor production in the LasR-RpoA* mutant is well in line with its QS-active phenotypes. To further assess the changes in bacterial virulence brought by LasR-RpoA*, we conducted cell-killing experiments using host cells of *P. aeruginosa*. Chinese hamster ovary (CHO) cells and human lung adenocarcinoma A549 cells were exposed to the test strains, and the induced cell death was quantified by measuring the release of cytosolic lactate dehydrogenase (LDH) from the cytosol. Consistent with its intermediate level of QS-dependent virulence factors, LasR-RpoA* caused an enhanced cell death compared to the LasR-null mutant, while demonstrated lower cytotoxicity than the LasR-MexT mutant (Fig. 7). Overall, our results show that the non-MexT mutant LasR-RpoA* enables QS-reprogramming, albeit with a trade-off in compromised bacterial pathogenesis on host cells.

**Figure 7.**
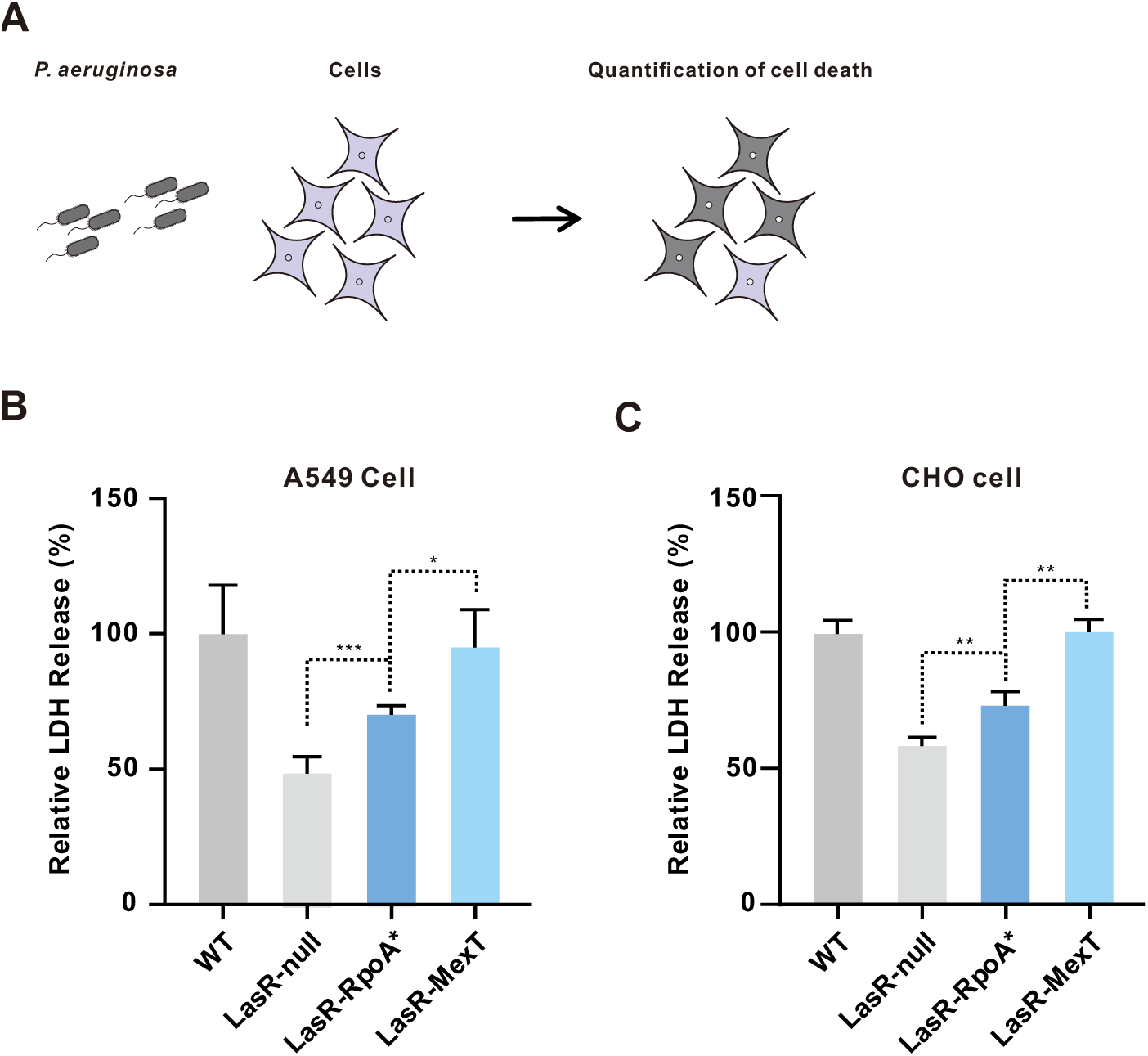
The RpoA* mutation significantly enhances host cell cytotoxicity. (A) Illustration of *P. aeruginosa* cell killing on host cells. (B-C) The killing assay with human lung cancer A549 cells (B) and Chinese hamster ovary (CHO) cells (C) was performed with equal amounts of the indicated strains. After incubation for 6 h, the release of cytosolic lactate dehydrogenase (LDH) from infected cells was quantified. The released amount of LDH inoculated with the wild-type strain (WT) was set to 100%. Data represent means ± SD (*n* = 3). \**P* < 0.05, ** *P* < 0.01, *** *P* < 0.001 (t-test).

## Discussion

A novel approach, named TGD-MS, was used in the evolution experiments of this study to discover mutations in genes involved in QS-reprogramming in a LasR-null mutant of *P. aeruginosa*. In the present study, the TGD-MS approach was used for obtaining QS-active revertants. By constructing a strain containing an extra copy of *mexT*, QS-active revertants carrying a mutated *mexT* gene could be filtered out to a large extent in our screen for QS-reprogramming mutants. Using this increased screening stringency, a QS-active mutant with a single nucleotide substitution in *rpoA* was identified. It is worth noting in this context that it would have been impossible to identify this new QS-reprogramming function of RpoA by conventional transposon insertional mutagenesis, because *rpoA* is an essential gene for cell viability (31). More generally, the TGD-MS method described in this work allows us to elucidate genetic components of known pathways and even to reveal unknown metabolic pathways. The use of the TGD-MS method in evolution experiments is now opening up new opportunities for scientists to make fundamental discoveries in a wide range of research areas. We hypothesize that the TGD-MS method can be employed to uncover genetic components of known and unexplored pathways in any transformable haploid organism.

Our study on a QS-inactive LasR-null mutant of *P. aeruginosa* uncovers a novel QS-reprogramming role for RpoA (Fig. 8A), which is an essential subunit of the bacterial RNAP. Adaptive mutations in *rpoA* have been found to be beneficial for bacterial growth in evolution experiments with different organisms (37, 38). Previous publications showed that mutations in *rpoA* of *Bacillus subtilis* and *Escherichia coli* promote the utilization of secondary carbon sources and ribosome synthesis (39). The QS-related function of RpoA* found in the present study implicates that the T262 to A substitution of this protein variant leads to a structural modification in RpoA which affects the interaction between RNAP and the promoter DNA of the *mexEF-oprN* operon (Fig. 5). These findings are in agreement with previous studies on bacteria that certain residues in the α-CTD domain of RpoA are important for promoter contact, while a subset of these residues is required for the interaction with transcriptional effectors (40–42). Overall, our results suggest that a minor structural change in RpoA caused by a single point mutation restores QS-dependent activities in a LasR-null mutant, thereby allowing adaptation to rapidly changing environments.

**Figure 8.**
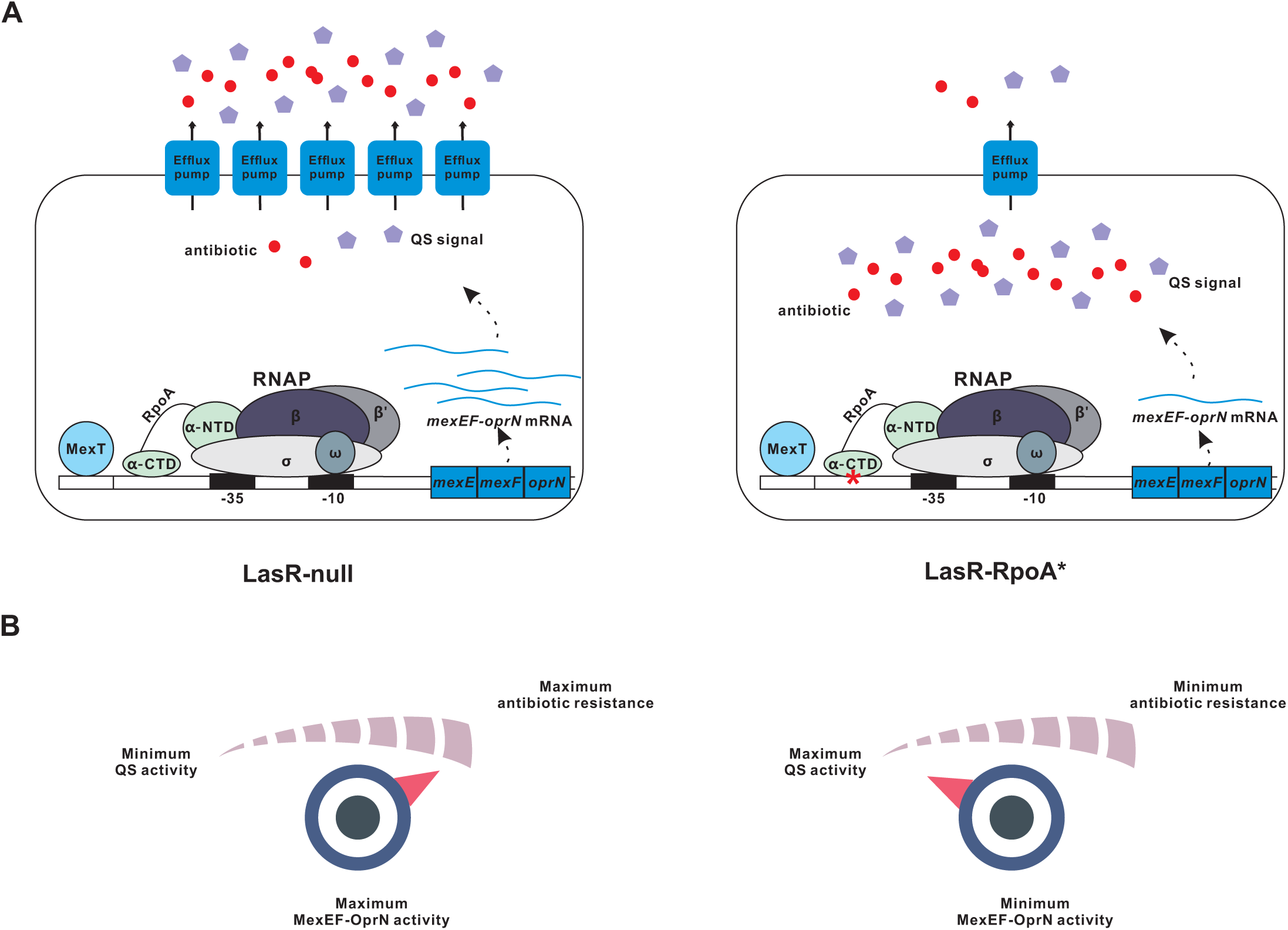
Schematic representation of the reduced activity of MexEF-OprN in LasR-RpoA*. (A) The LasR-RpoA* mutant induces down-regulated expression of *mexEF-oprN* operon genes, resulting in the reduction of intercellular exportation of antibiotics and QS signal molecules. Thus, the accumulation of antibiotics and QS signal confers bacterial antibiotic susceptibility, while activating the QS circuit. (B) The activity of the MexEF-OprN efflux pump positively correlates with antibiotic resistance while negatively associates with QS activity. For example, the activation of MexEF-OprN enhances antibiotic resistance, but decreases QS activity and vice versa.

LasR-MexT mutants of *P. aeruginosa* exhibit low expression of the *mexEF-oprN* operon, suggesting that weak or absent protein levels of the MexEF-OprN efflux pump lead to QS-reprogramming in evolution experiments (13, 14). Similarly, the QS-active LasR-RpoA* mutant in our study showed QS-related activities and transcriptome changes, including reduced expression of *mexEF-oprN* operon genes. Our transcriptome analysis, EMSA and the QS-active phenotype of the LasR-RpoA* mutant collectively suggest that RpoA* affects the QS network through expression of *mexEF-oprN* operon genes. The MexEF-OprN efflux pump exports the PQS precursor HHQ and inactivation of this pump is therefore increases PQS levels within the cell, resulting in induction of PQS-related QS responses (17, 33). However, deletion of key components of the PQS system, such as PqsA and/or PqsE (6, 34), only partially reduced QS activity in a LasR-MexT mutant (13). Similarly, the LasR-RpoA*-PqsA mutant deficient in PQS synthesis was found to possess weak but detectable QS activity in the present study, indicating that the RpoA*-mediated QS activation in LasR-RpoA* is partially independent of the PQS system.

Multidrug-resistance pumps of the RND family play key roles in bacterial resistance against antibiotics (43). The *mexEF-oprN* operon of *P. aeruginosa* codes for a typical RND efflux pump, which renders the bacterium largely resistant to certain antibiotics, such as chloramphenicol, fluoroquinolones and trimethoprim (18). On the other hand, the MexEF-OprN efflux pump can also export HHQ, and overexpression of *mexEF-oprN* operon genes resulted in reduced QS-related activities (17, 33). Thus, protein levels of this efflux pump may ultimately affect QS-controlled bacterial virulence of clinical *P. aeruginosa* isolates. Previous studies reported that CF patients possess QS-active LasR-null strains in their lungs (8, 10). *P. aeruginosa* isolates from chronically infected CF patients often develop antibiotic resistance because they receive continuous high doses of antibiotics over a long period of time (44). In other words, the expression of efflux pump genes is required for antibiotic resistance but impairs QS circuits that control the production of bacterial virulence factors. This raises the question of how QS-inactive LasR-null mutants resolve the dilemma between antibiotic resistance and QS-reprogramming. The findings of this study highlight that the MexEF-OprN efflux pump is at the center of this tradeoff between antibiotic resistance and QS-controlled virulence factor production (Fig. 8B). The single amino acid substitution in RpoA* caused reduced expression of the *mexEF-oprN* operon genes in our study. Accordingly, when compared with the LasR-MexT mutant, the LasR-RpoA* mutant was found to have relatively higher growth fitness in competing with the LasR-null parental strain when the bacteria were exposed to increasing concentrations of chloramphenicol. In conclusion, our work shows that a mutation in *rpoA* can fine-tune the efficiency of an RND efflux pump required for antibiotic resistance and efflux of HHQ, thereby modulating antibiotic resistance and the production of QS-dependent virulence factors under varying environmental conditions.

*P. aeruginosa* possesses a complicated regulatory gene network to control the expression of virulence genes (30) and undergoes evolutionary changes in the phase of chronic CF infection (44). In our TGD-MS approach for example, when an extra copy of *mexT* was inserted into a LasR-null mutant, we observed that the bacterium can activate MexT-independent alternative QS-reprogramming pathways. Given that MexT controls not only the expression of *mexEF-oprN* operon genes but also the transcription of potentially important virulence genes (20, 21), it can be speculated that maintaining a non-modified MexT is critical for survival in chronically infected CF patients. Notably, our *mexT* sequence analysis and previous observations indicate that numerous LasR-null, QS-active *P. aeruginosa* isolates from CF patients retain a functional MexT (9, 15). Taken together, we report in this study on a novel experimental evolution approach that facilitated the identification of non-*mexT* genes involved in the formation of the MexEF-OprN efflux pump and QS-reprogramming. Future research is needed to identify similar mutations in clinical isolates and to clarify their effects on QS activities, virulence and antibiotic resistance.

## Materials and methods

### Bacterial strains and growth conditions

*P. aeruginosa* PAO1 and mutant derivatives (Supplementary Table S6) were grown in Luria Bertani (LB) broth (containing 10 mg/ml tryptone, 5 mg/ml yeast extract, 10 mg/ml NaCl) at 37°C. LB broth was buffered with 50 mM 3-(N-morpholino) propanesulfonic acid, pH 7.0 (LB-Mops broth). The photosynthetic medium (PM) (45) supplemented with 1% sodium caseinate (Sigma Aldrich, St. Louis, MO, USA) as the sole carbon source (PM-casein medium) was used for evolution experiments at 37°C. Unless otherwise specified, *P. aeruginosa* strains were cultured in 14-mL Falcon tubes (Corning Inc., Corning, NY, USA) containing 3 ml medium, with shaking (220 rpm) at 37°C. *Escherichia coli* was grown in LB broth at 37°C. Details on bacterial strains and plasmids used in this study are shown in Supplementary Table S6.

### Construction of *P. aeruginosa* mutants

Construction of *P. aeruginosa* mutants (LasR-MexT with *mexT* deletion, LsR-RpoA* mutant with a *rpoA* point mutation (T262→A), LasR-RpoA*-PqsA with a *pqsA* deletion) was performed according to a homologous recombination approach described previously (46). Briefly, about 500 ∼ 1000 bp of DNA flanking the targeted single nucleotide substitution or the full length gene of interest were PCR-amplified and cloned into the pEXG-2 vector containing a gentamycin (Gm) resistance gene (46, 47) using the Vazyme ClonExpress II One Step Cloning Kit (Vazyme Biotech, Nanjing, China). The constructed plasmids were then mobilized into *P. aeruginosa* (LasR-null and LasR-RpoA*, respectively) by triparental mating with the help of *E. coli* carrying the helper plasmid pRK2013 (Figurski and Helinski, 1979). Mutants were first selected on *Pseudomonas* Isolation Agar (PIA) containing Peptic digest of animal tissue 20.0 g/L,Magnesium chloride 1.4 g/L, Potassium sulfate 10.0 g/L, Triclosan (Irgasan) 0.025 g/L, and Agar 13.6 g/L supplemented with 100 μg/ml Gm and then on LB agar containing 10% (w/v) sucrose. All mutants were confirmed by PCR amplification and subsequent DNA Sanger sequencing. The primers used in this study are listed in Supplementary Table S7.

### Complementation of the LasR-RpoA* mutant

The *rpoA* gene was cloned into the pJN105 expression vector with the Vazyme ClonExpress II One Step Cloning Kit, generating plasmid pJN105-*rpoA*. The constructed plasmid and the empty vector pJN105 as a control were then mobilized into the LasR-RpoA* mutant by triparental mating. The bacteria were grown in LB-Mops broth at 37°C. Expression of *rpoA* in the complementation strain was induced by addition of 1 mM L-arabinose and the bacteria were harvested 20 h later.

### Evolution experiments

Three independent colonies of the LasR-null mutant carrying a miniTn7-MexT construct at a neutral site of the genome were separately inoculated into 3 ml LB-Mops broth to obtain an overnight culture. 100 μL of the bacterial suspension were then transferred into 3 ml PM-casein medium in 14-mL Falcon tubes (Corning). One passage was performed every 5 days by transferring 100 μL of bacterial suspension into 3 ml fresh PM-casein medium. When required, water was added to compensate for evaporated medium. The bacterial suspensions were then spread on LB agar plates. Finally, to visualize extracellular protease activity of the obtained colonies, the bacteria were subjected to a skim milk assay. An initial evolution experiment with a non-modified LasR-null mutant population (carrying a single *mexT* gene copy) was performed in a similar way. Furthermore, an iterative evolution experiment was performed with a LasR-null mutant carrying miniTn7-*mexT* and an additional miniTn7-*rpoA* construct.

### Skim milk assay

Total extracellular proteolytic activity of *P. aeruginosa* strains was evaluated using skim milk agar plates on which the bacteria form a protease-catalyzed clearing zone surrounding each colony. Individual colonies were grown on LB agar and then spotted on the skim milk agar plates (25% (v/v) LB, 4% (w/v) skim milk, 1.5% (w/v) agar). Colonies with extracellular proteolytic activity formed clearing zones after incubation at 37°C for 24 h. The area of transparent zones reflecting extracellular proteolytic activity was quantified from captured photographs.

### Pyocyanin measurement

Overnight cultures of *P. aeruginosa* grown in LB-Mops broth were diluted into 4 ml *Pseudomonas* P broth (20 g/L pancreatic digest of gelatin, 1.4 g/L magnesium chloride, 10 g/L potassium sulfate) to reach a starting OD_600_≈0.02. The bacteria were cultured at 37°C for 20 h. The cells were then centrifuged at 16,000 g × 2 min. The culture supernatants were collected and their OD values at 695 nm were photometrically determined at 695 nm. Pyocyanin production was estimated by determining OD_695_/OD_600_ values over time.

### Hydrogen cyanide measurement

A test paper method was used to detect hydrogen cyanide produced by *P. aeruginosa* strains. A circle of a Whatman 3MM chromatography paper corresponding to the size of an agar plate was soaked in the HCN detection reagent consisting of 5 mg copper (II) ethyl acetoacetate 5 mg 4,4’-methylenebis-(N, N-dimethylaniline) mixed with chloroform (1 to 2 ml). *P. aeruginosa* cells grown overnight in LB-Mops broth were used to spot-inoculate 2% (w/v) peptone agar plates. After incubation at 37°C for 12 h, the plates were overlaid with the prepared test paper and incubated at 37°C for additional 12 h. The color of the paper turned blue when it was exposed to hydrogen cyanide. Image J software (https://imagej.nih.gov/ij) was used to quantify the production of hydrogen cyanide.

### Swarming motility assay

*P. aeruginosa* strains were cultured in LB-Mops broth overnight to reach an OD_600_≈1.8. Each bacterial suspension (1.5 μL; OD_600_ adjusted to 0.5) was then transferred to the center of an agar plate containing 0.5% (w/v) Bactor-agar, 8g/L nutrient broth (BD, CA, USA) and 5 g/L glucose. Bacteria were incubated at 37°C for 16-18 h.

### Labeling of miniTn7-Gm

The pUC18T-mini-Tn7T-Gm (NCBI accession number: AY599232) (48)was integrated into a neutral site of the PAO1 genome by triparental mating using the helper plasmid pTNS2 (NCBI accession number: AY884833). The integration event was confirmed by PCR amplification and DNA sequencing. The excision of the Gm resistance gene was performed with the pFLP2 plasmid (NCBI accession number: AF048702) (48) and selection on LB agar containing 5% (w/v) sucrose.

### Assays with strains containing reporter plasmids

Plasmids with a *rhl*Α promoter*-*GFP fusion (8) and a constructed *pqs*Α promoter-GFP fusion were used to quantify the RhlR- and PQS-responsive activities, respectively. A plasmid containing a *mexE* promoter-GFP construct was used to assess the expression of the *mexEF-oprN* operon. The reporter plasmids were mobilized into *P. aeruginosa* strains and selected on LB agar plates (containing Gm). PAO1 strains carrying QS reporter plasmids were grown in LB-Mops broth containing 50 mg/ml Gm for 12 h. The bacterial suspensions were then diluted to LB-Mops broth containing 50 mg/ml Gm (OD_600_ ≈ 0.02) and grown for 18 h to stationary phase. Finally, the bacteria were transferred to 96-well plates (200 µl/well) with three technical replicates. Fluorescence (excitation 488 nm, emission 525 nm) and optical density (OD_600_) values of the samples were determined using a microplate reader (Synergy H1MF, BioTek Instruments, Winooski VT, USA).

### Quantification of C4-HSL and PQS

*P. aeruginosa* strains were cultured overnight in LB-Mops broth. Bacteria were then grown in 4 ml LB broth at 37°C for 18 h (starting OD_600_≈0.02). AHLs were extracted with an equal amount of ethyl acetate. A bioassay with the reporter *E. coli* strain containing the pECP61.5 with a *rhlA-lacZ* fusion and an IPTG-inducible *rhlR* (8) was used to quantify C4-HSL in the ethyl acetate extract. Precise quantification of PQS was performed by LC/MS as described previously (34). 10 μl of the ethyl acetate phase was subjected to liquid chromatography-mass spectrometry (LC-MS) analysis. The detection system (Q-Exactive Focus/Ultimate 3000; Thermo Fisher Scientific, Waltham, MA, USA) was equipped with a 100 x 2.1 mm, 1.7 µm ACQUITY UPLC HSS T3 chromatographic column (Waters, Milford, MA, USA). An acidified (1% glacial acetic acid by volume) water/methanol gradient was used as the mobile phase (0.4 ml/min flow rate; 40°C). PQS was quantified by measuring the area of PQS peaks in chromatograms from different samples and values were standardized according to the concentration of an added internal standard.

### Production and purification of RpoA proteins

Full-length of His-tagged RpoA or the RpoA* variant was cloned into pJN105 expression vector. The plasmid was then mobilized into the LasR-RpoA* mutant and protein expression was induced by growth in LB broth containing 1 mM L-arabinose at 37°C for 18 h. Cells were pelleted at 13,000 g × 5 min and resuspended in lysis buffer (PBS buffer supplemented with 1 × Protease Inhibitor Cocktail (Bimake, Houston, TX, USA), 1mg/ml lysozyme). Resuspended cells were lysed in a high pressure cell homogenizer machine (Stansted Fluid Power Ltd, Essex, United Kingdom) at 420 MPa. The lysed cells were centrifuged at 13,000 g × 5 min (4°C). The supernatant was then incubated with Ni-NTA beads (Qiagene, Shanghai, China) at 4°C for 2 h. The mixture was loaded on a chromatographic column, washed with washing buffer (500 mM NaCl, 20 mM NaH_2_PO_4_, 20 mM imidazole, pH 8.0), and finally eluted with a buffer containing high concentrations of imidazole (500 mM NaCl, 20 mM NaH_2_PO_4_, 250 mM imidazole, pH 8.0). Collected fractions were assessed by Native PAGE analysis. Fractions containing recombinant protein were pooled and concentrated with an ultrafiltration device (MilliporeSigma, Burlington, MA, USA) and finally eluted with PBS buffer. Samples were stored at -80°C.

### Electrophoretic mobility shift assay (EMSA)

The promoter sequence of the *mexEF-oprN* operon (P*mexE*) was used as probe for EMSA. Probe DNA was labelled according to our published method (49). Briefly, the probe DNA was labeled with Biotin-11-UTP (Jena Bioscience, Jena, Germany) through PCR amplification using T4 DNA polymerase (Novoprotein, Suzhou, China). The labeled probe (30 ng) was incubated with 0.2 μg purified His-tagged RpoA or RpoA* protein in binding buffer (10 mM Tris-HCl, 50 mM KCl, 1 mM DTT, 1% (v/v) glycerin, 10 mM MgCl_2_, 1 μg/μL poly(deoxyinosinic-deoxycytidylic) acid, pH 7.5) at room temperature for 30 min. The samples containing a given DNΑ-protein complex were subjected to electrophoresis on a 5% polyacrylamide gel in 0.5×Tris-borate-EDTA (TBE) buffer at 100 V for 180 min. DNA subsequently was transferred to a nylon membrane and incubated with milk blocking buffer (50 g/L milk powder in TBST buffer (50 mM Tris, 150 mM NaCl, 0.05% Tween-20, pH 7.5) for 30 min with shaking at room temperature. The membrane was then incubated with streptavidin-horseradish peroxidase conjugate (Thermo Fisher Scientific) in blocking buffer for 30 min at room temperature. The membrane was subsequently washed with TBST buffer for 4 times, 5 min each. DNA was visualized using the Immobilon Western kit (Millipore) and photographed by a Tanon 5200 Multi-Imaging System (Tanon, Shanghai, China).

### Mammalian cell cytotoxicity assay

Human lung cancer A549 cells and Chinese hamster ovary (CHO) cells were cultured in DMEM medium (GIBCO, Thermo Fisher Scientific, China) and RPMI1640 medium (GIBCO, Thermo Fisher Scientific, China) supplemented 10% FBS (GIBCO), respectively, at 5% CO_2_ and 37°C. Exponentially growing *P. aeruginosa* bacteria cultured in LB broth (OD_600_≈0.6) were collected by centrifugation (4,000 g × 5 min) and washed with PBS. Next, the cells were added to near-confluent CHO and A549 cells at a starting multiplicity of infection (MOI) ratio of 5: 1. After incubation at 37°C for 6 h, the extent of cell killing was determined by quantification of the release of lactate dehydrogenase into the cell culture supernatant using the LDH Cytotoxicity Detection Kit (Beyotime, Nantong, China).

### Competition experiment

The *P. aeruginosa* strains (LasR-null, LasR-RpoA* and LasR-MexT) were grown in PM medium supplemented with 0.5% (w/v) casein amino acids (Sangon, Shanghai, China) overnight and then transferred to liquid PM-Casein medium (adjusted to OD_600_≈0.02) with or without chloramphenicol. Each strain tagged with a pUC18T-mini-Tn7T-Gm construct was cocultured with the non-tagged strain at a starting ratio of 5:95. The mixed cells were then grown at 37°C for 48 h. Cell counts were determined by spreading the bacteria onto LB agar plates supplemented with or without 10 μg/ml Gm. The relative fitness of each strain was deduced from the ratio of Malthusian growth parameters (*w*) = ln(*X*_1_/*X*_0_)/ln(*Y*_1_/*Y*_0_), as defined previously (50).

### Extraction and sequencing of RNA

*P. aeruginosa* strains were grown in LB-Mops broth at 37°C for 18 h (OD_600_ ≈ 3.8-4.0). Total RNA was isolated using TRIzol™ Reagent (Thermo Fisher Scientific). The quantity of extracted total RNA was measured using a NanoDropTM 1000 spectrophotometer (Thermo Fisher Scientific). Extracted RNA was treated with DNase (RQ1 RNase-free DNase I, Promega, Beijing, China) to remove traces of genomic DNA. The obtained total RNA (2 independent RNA samples per strain) was sent to Novagen (Tianjin, China) for stranded paired-end mRNA-seq sequencing using the Illumina Novaseq Platform.

### Quantitative real-time PCR (qRT-PCR)

Total RNA of *P. aeruginosa* strains was reverse transcribed using the HiScript II 1st Strand cDNA Synthesis Kit (Vazyme Biotech) following the manufacturer’s instructions. The obtained cDNA was then used for qRT-PCR analysis. Reactions containing the SYBR qPCR Master Mix (Vazyme Biotech) were prepared in 96-well plates and run in a StepOnePlus Real-Time PCR System (Applied Biosystems, Waltham, MA, USA) as recommended by the supplier. Primers used for qRT-PCR are listed in Table S7. The measured values for a given gene were normalized to the expression level of the *proC* gene. Reactions were performed in triplicate.

### Transcriptome analysis

*P. aeruginosa* strains (LasR-null, LasR-RpoA* and LasR-MexT, each strain has two replicates) were subjected to RNA-seq analysis. RNA-seq short reads were filtered with a customized Perl script to remove Illumina adapters and reads of low quality (Phred Score, Q < 30). The quality of filtered reads was assessed by FastQC. Trimmed reads were then mapped to the *P. aeruginosa* PAO1 reference genome (NC_002516.2) using Hisat2. The SAM files were further converted to BAM files by Samtools. Reads aligned to the reference genome were counted by HTSeq. The generated count files were further processed with the R package DESeq2. Genes were regarded as “differentially expressed” if they showed a fold change ≥2.0 and a *P* value ≤0.05.

### Gene expression profile analysis

Genes showing 4-fold expression changes in strain comparisons were compared using customized R and Perl scripts. *P* values were computed using a one-sided KS test (51).

### Used Software

The following software were used in this study:

BWA software, version 0.7.15-r1140 (http://bio-bwa.sourceforge.net) (52);

Cutadapt software, version 1.16 (https://github.com/mafdrcelm/cutadapt/) (53);

Samtools software, version 1.5 (http://samtools.sourceforge.net)(54);

FastQC, version fastqc_v0.11.5 (https://www.bioinformatics.babraham.ac.uk/projects/fastqc/);

Hisat2, version 2.1.0 (https://daehwankimlab.github.io/hisat2/) (55);

HTSeq, version 0.11.1 (https://htseq.readthedocs.io/en/release_0.11.1/count.html) (56);

DESeq2 (57);

Perl software, version v5.22.1 (https://www.perl.org/);

Python software, version v3.8.2 (https://www.python.org/);

R software, version v3.6.1 (http://www.R-project.org/);

GraphPad Prism software, version 5 (https://www.graphpad.com/);

Image J software, version 1.8.0_172 (http://imagej.nih.gov/ij/).

### Statistical analysis

Statistical analysis (*t*-test and KS tests) was performed using Excel and R software (http://www.R-project.org/).

### Data Availability

RNA-seq data sets have been deposited in the NCBI SRA database (www.ncbi.nlm.nih.gov/sra) under the accession number PRJNA863086 and PRJNA962310.

## Authors’ contributions

W.C., H.L., M.L., X.Z., X.C. and W.D. designed experiments. W.C., H.L., M.L., X.Z., X.C. and W.D. conducted the experiments. W.D. performed bioinformatics analyses. W.C., W.D. and C.S. wrote the manuscript. All authors reviewed the manuscript.

## Acknowledgements

We are grateful to Eugene Makeyev (King’s College London, UK) for providing useful assistances in bioinformatics analyses. We thank E. Peter Greenberg (University of Washington, USA) kindly donated the *P. aeruginosa* wild-type strain PAO1. We want to extend our gratitude to all lab members for their assistances and supports with respect to many aspects of this work.

