## Supplemental Figure S1 to S7 for "Discovery of RNA polymerase α-subunit protein as a novel quorum sensing reprograming factor in *Pseudomonas aeruginosa*"

**Figure S1.** The constructed LasR-RpoA* mutant exhibits elevated proteolytic activity.

**Figure S2.** Complementation of the LasR-RpoA* mutant with a wild-type *rpoA* gene reduces QS-related activities.

**Figure S3.** qRT-PCR validation of differentially expressed genes obtained from the transcriptomic analysis.

**Figure S4.** Deletion of *pqsA* in LasR-RpoA* reduces RpoA*-mediated extracellular protease activity.

**Figure S5.** Complementation of the LasR-RpoA* mutant with a copy of *mexEF-oprN* reduces pyocyanin production.

**Figure S6.** Swarming motility in the LasR-RpoA* mutant.

**Figure S7.** The production of QS-controlled metabolites in the LasR-RpoA* mutant.

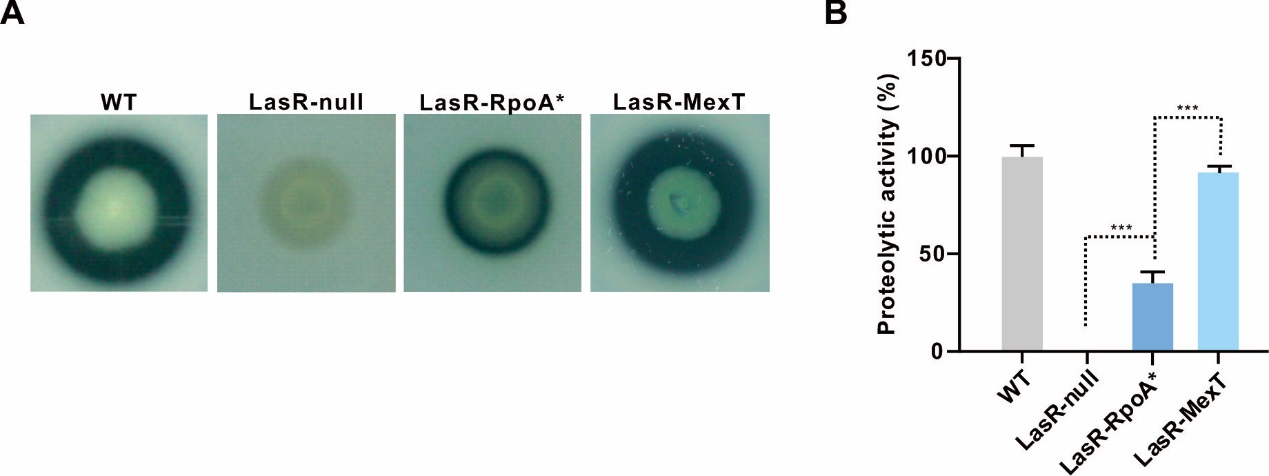

**Figure S1. The constructed LasR-RpoA* mutant exhibits elevated proteolytic activity.**

(A) Equal numbers of bacteria of the indicated strains were spotted on skim milk plates to estimate proteolytic activity resulting in transparent proteolytic zones (halos). Pictures were taken 24 h later. (B) Quantification of proteolytic zones shown (A). Data represent means ± SD (*n* = 3). **P* < 0.05, ***P* < 0.01, ****P* < 0.001 (*t*-test).

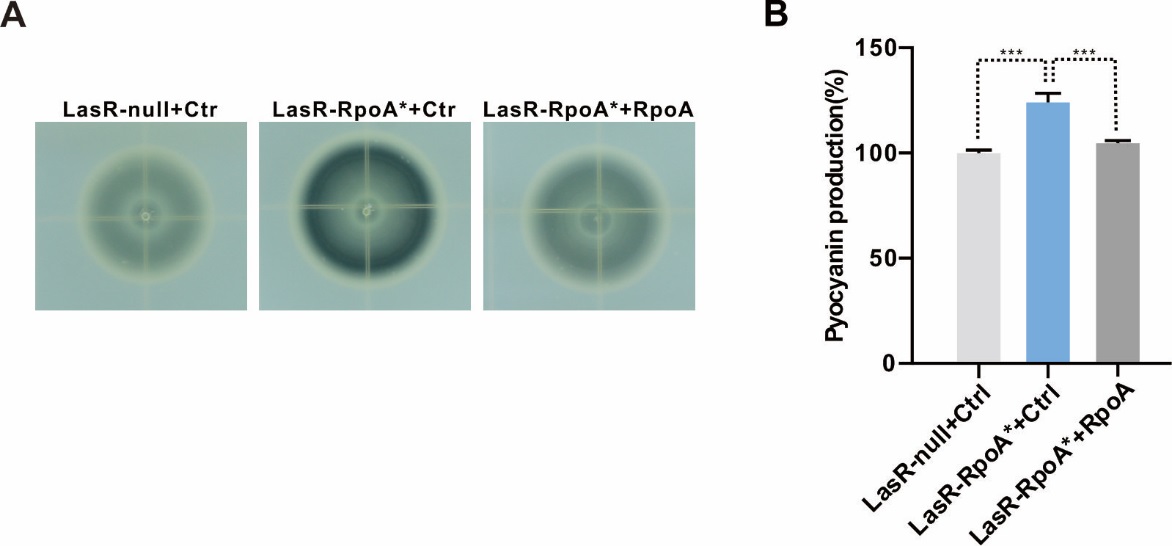

**Figure S2. Complementation of the LasR-RpoA* mutant with a wild-type *rpoA* gene reduces QS-related activities.**

Introduction of an episomal copy of the wild-type *rpoA* gene into the LasR-RpoA* mutant diminished the QS-controlled proteolytic activity and pyocyanin production. (A) Proteolytic activity as visualized by the skim milk plate assay. Photographs of indicated strains were taken after the transfer of equal amounts of bacteria onto the skim milk plate and incubation at 37^o^C for 24 h. (B) Pyocyanin production of indicated strains. The LasR-null mutant or the LasR-RpoA* mutant was either transformed with the empty vector pJN105 (Ctr) or the RpoA-expressing vector pJN105-RpoA (RpoA). Expression of *rpoA* was induced by the addition of 1 mg/ml L-arabinose. Data represent means ± SD (*n* = 3). **P* < 0.05, ***P* < 0.01, ****P* < 0.001 (*t*-test).

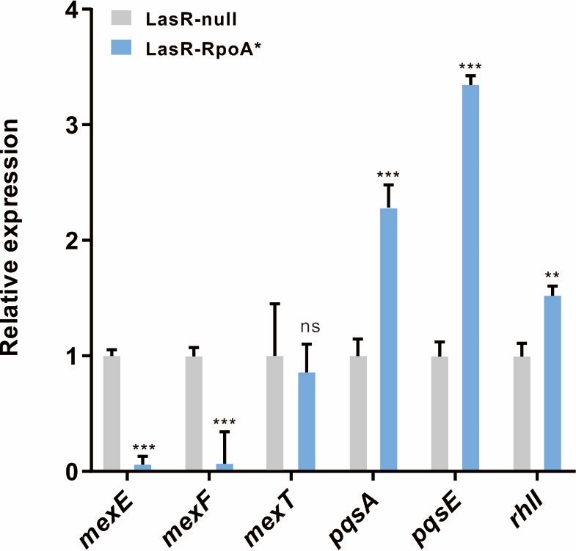

**Figure S3. qRT-PCR validation of differentially expressed genes obtained from the transcriptomic analysis.**

The qRT-PCR validation of differentially expressed genes obtained from the transcriptome analysis. Relative expression was normalized using *proC* gene data. Data represent means ± SD (3 independent RNA extractions; *n* = 3). **P* < 0.05, ***P* < 0.01, ****P* < 0.001 (*t*-test).

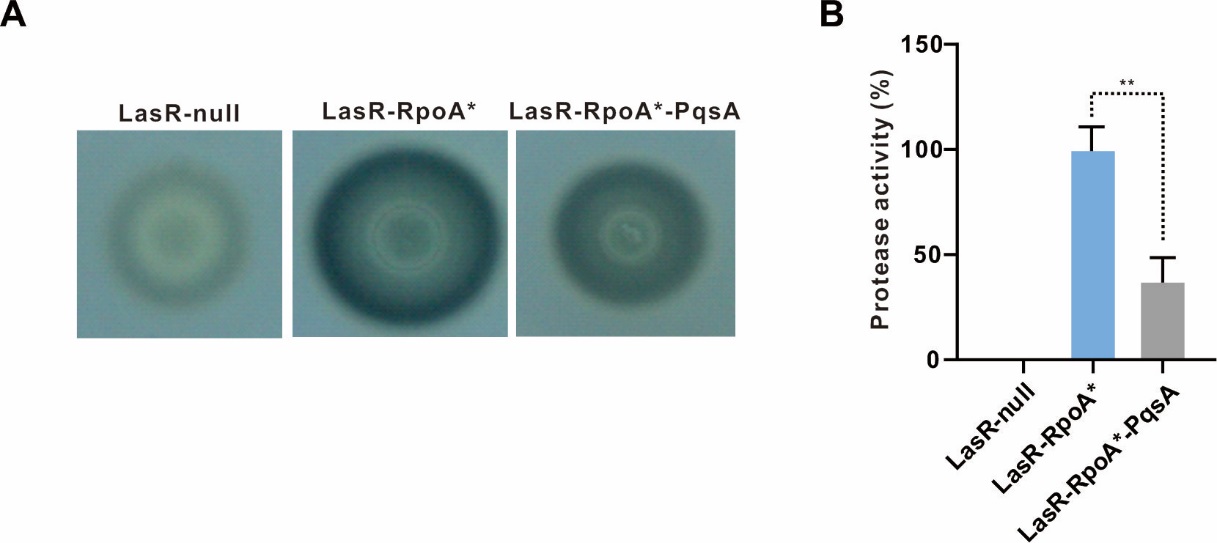

**Figure S4. Deletion of *pqsA* in LasR-RpoA* reduces RpoA*-mediated extracellular protease activity.**

(A) Extracellular proteolytic activity was visualized through spotting equal amounts of indicated strains on skim milk agar plates. Plates were photographed after incubation at 37^o^C for 24 h and representative photos are shown. (B) Quantification of proteolytic zones obtained from obtained pictures. Data represent means ± SD (*n* = 3). **P* < 0.05, ***P* < 0.01, ****P* < 0.001 (*t*-test).

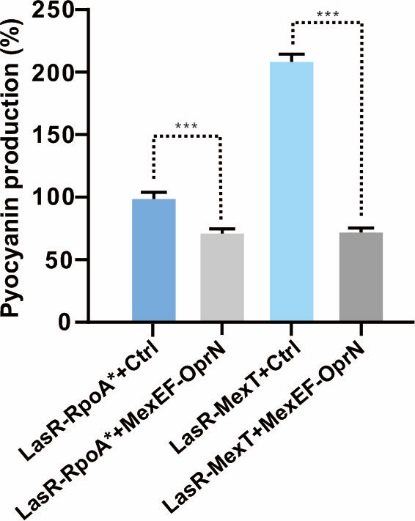

**Figure S5. Complementation of the LasR-RpoA* mutant with a copy of *mexEF-oprN* reduces pyocyanin production.**

The LasR-RpoA* was complemented by introducing a miniTn7-based construct containing the *mexEF-oprN* operon driven by a *rrnB* promoter. Pyocyanin production (OD_695_/OD_600_ values) was measured in indicated strains (+Ctr, mutant strain carrying the empty miniTn7 vector; +MexEF-OprN, the mutant strain complemented with miniTn7 containing wild-type *mexEF-oprN* driven by an *rrnB* promoter). Data represent means ± SD (*n* = 3). **P* < 0.05, ***P* < 0.01, ****P* < 0.001 (*t*-test).

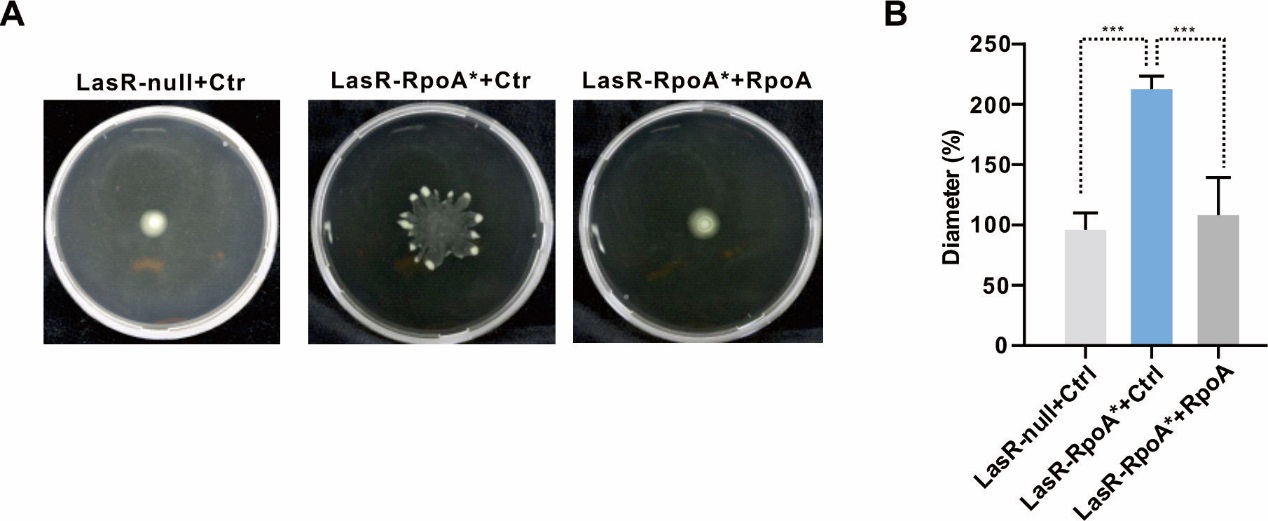

**Figure S6. Swarming motility in the LasR-RpoA* mutant.**

(A) Bacterial swarming motility was assayed on 0.5% agar plates. Plates were photographed after incubation at 37^o^C for 18 h. Representative photographs are shown for indicated mutant strains (LasR-null-Ctr, the LasR-null mutant carrying the empty pJN105 vector; LasR-RpoA*-Ctr, the LasR-RpoA* mutant carrying the empty pJN105 vector; LasR-RpoA*-RpoA, the LasR-RpoA* mutant complemented with pJN105-RpoA containing wild-type *rpoA*). (B) Quantification of diameters in formed colonies. The *rpoA* gene expression was induced by the addition of L-arabinose (1 mg/ml). Data indicate means ± SD from a representative experiment (*n* = 3). *P* values were obtained from *t*-tests. **P* < 0.05, ***P* < 0.01, ****P* < 0.001.

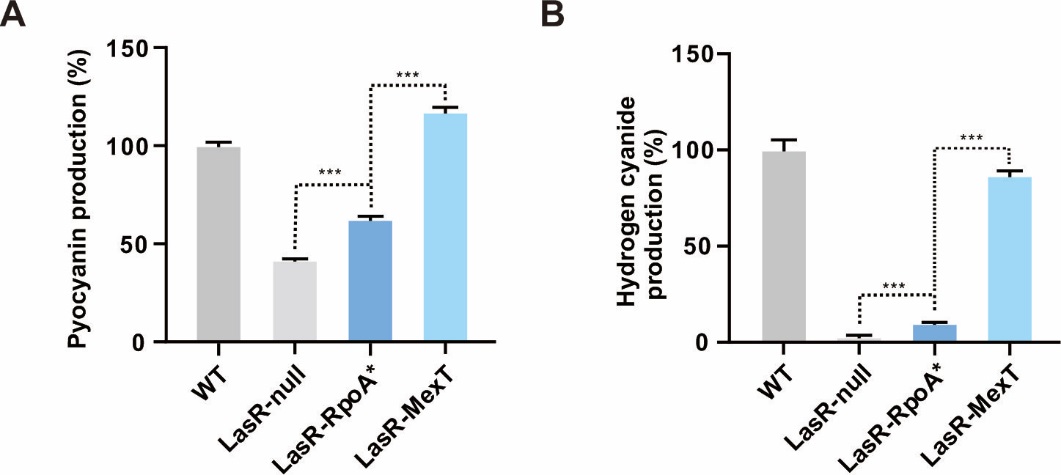

**Figure S7. The production of QS-controlled metabolites in the LasR-RpoA* mutant.**

(A) Pyocyanin production (OD_695_/OD_600_ values) in shown strains. (B) Cyanide production by these strains. Image J software was used to compare the amount of cyanide produced. The cyanide-sensitive filter papers were photographed after growth of the bacteria in 6-well plates at 37^o^C for 24 h. The amount of cyanide produced in wild-type PAO1 was set to 100%. Data are presented as means ± SD (*n* = 3, *t*-test). **P* < 0.05, ***P* < 0.01, ****P* < 0.001 (*t*-test).
